# Early spiral arteriole remodeling in the uterine-placental interface: a rat model

**DOI:** 10.1101/2023.08.13.553146

**Authors:** Sarah J. Bacon, Yuxi Zhu, Priyanjali Ghosh

## Abstract

The mammalian placenta’s interface with the parent is a richly vascularized tissue whose development relies upon communication between many different cell types within the uterine microenvironment. The uterine blood vessels of the interface are reshaped during pregnancy into wide-bore, flaccid vessels that convey parental blood to the exchange region of the placenta. Fetally-derived invasive trophoblast as well as parental uterine macrophages and Natural Killer cells are involved in the stepwise remodeling of these vessels and their respective contributions to this crucial process are still being delineated. However, the earliest steps in arteriole remodeling are understudied as they are difficult to study in humans, and other species lack the deep trophoblast invasion that is so prominent a feature of placentation in humans. Here, we establish the rat, with deep hemochorial placentation akin to humans, as a model system in which to tease apart the earliest, relatively understudied events in spiral arteriole remodeling. We show that the rat uterine placental interface increases in size and vascularity rapidly, before trophoblast invasion. The remodeling stages in the arterioles of the rat uterine-placental interface follow a sequence of anatomical changes similar to those in humans, and there are changes to the arterioles’ muscular tunica media prior to the marked influx of immune cells. The rat is a tractable model in which to better understand the cell/cell interactions occurring in vivo in an intact tissue microenvironment over time.

## Introduction

In healthy pregnancy, fine uterine arterioles reaching into the deep placental bed are widened into large-radius tubes that deliver blood to the placenta’s intervillous space. A regulated remodeling process increases the vessel bore, lowering the velocity, pulsatility, and pressure at which blood flows into the placental sinus where delicate villous trees coated with fetal trophoblast are suspended (1). Vessel remodeling that is incomplete or that fails to reach into the myometrial segment of the uterine arterioles has been associated with preeclampsia (2), fetal growth restriction (3,4) and placental abruption (5). Healthy placental development is plastic, matching fetal needs to parental capacity in the face of variation in oxygen or nutrient availability (6,7), with profound effects on not just prenatal but also postnatal development. Placental development and the remodeling of uterine vessels in the deep placental bed are thus crucial to both healthy pregnancy and postnatal life.

Remodeling of these convoluted uterine arterioles, or “spiral arteries,” is a feature of both human and rodent pregnancy. By 16-18 weeks’ gestation in humans (8), and by 18 days in rat (9), parental spiral arteries in the deep placental bed have been transformed into large-radius, low-resistance vessels. The muscular, elastic tunica media layer of the arteriole becomes disorganized and fragmented, releasing the vessel from parental vasomotor control over radius. Remodeled vessels dilate to several times their original size, and the vessel’s vasoactive tunica media is ultimately replaced by trophoblast embedded in a fibrinoid matrix (10). The cellular mechanisms responsible for this choreographed vessel deconstruction and rebuilding are of keen research interest.

Trophoblast is an invasive cell type originating from the outer trophectoderm of the blastocyst. These cells break off the placenta’s chorionic plate and migrate into the uterine parenchyma, where they associate with the parental spiral arterioles. Two invasive trophoblast lineages develop, interstitial and endovascular (11,12). Interstitial trophoblast cells migrate into the stroma around vessels in the deep placental bed (13), and endovascular trophoblast migrates from the placenta into the arterioles themselves in humans and rodents (9), ultimately replacing the muscular media layer of the arteriole (14,15,13) and rendering these vessels chimeric.

The physiological change of a vessel into a sac-like, dilated vessel capable of high flow (16,17) is termed “trophoblast-dependent” remodeling. Indeed trophoblast can induce phenotypic changes in vascular smooth muscle consistent with a role in this transformation. Human trophoblast can induce muscle cell apoptosis in vessel explants (18), degrade vessel elastin (19), and stimulate vascular smooth muscle to de-differentiate *in vitro* (20). In rats, the presence of extravillous trophoblast is correlated with the phenotypic switching of vascular smooth muscle from a contractile to a proliferative and noncontractile form (21).

Yet many anatomical and functional changes to the uterine vasculature precede trophoblast invasion. Human spiral arterioles dilate and the muscular tunica media layer disorganizes (22) and loses alpha actin activity prior to trophoblast invasion (23), (24). In rats, trophoblast only enters the vascular compartment on day 13 of gestation (7,9) yet on days 11 and 12 uterine blood flow has begun to shift away from the parietal decidua and toward the chorioallantoic placenta (25), suggesting that in rats at least, vascular resistance has dropped in the vessels of the deep placental bed prior to the invasion of trophoblast. It seems that trophoblast-dependent changes are not a complete explanation for vessel remodeling.

This earlier, potentially preparatory phase of vessel remodeling is often termed “decidua-associated” remodeling (26,27) as it depends on cells and signals within the decidual microenvironment. The loss of vascular media integrity in this early phase remodeling is coincident with the invasion of maternal leukocytes in humans and in mice, including both uterine Natural Killer cells and macrophages (28–31). Uterine Natural Killer cells are especially important: in culture, uNKs initiate vascular smooth muscle cell rounding, layer separation, and misalignment, effects blocked by protease inhibitors (32). Depleting uNKs in mice results in a failure of vessel remodeling (33–35).

But trophoblast and leukocyte activity are intertwined, suggesting that compartmentalizing the remodeling process into two discrete phases may be oversimplified. Perhaps most significantly, uNK cells influence trophoblast differentiation in humans (36) and in rat regulate trophoblast lineage determination (7) through their release of proangiogenic factors that influence oxygen delivery to the trophoblast. Depletion of uNK in rat or mouse induces a more invasive trophoblast phenotype (34,37) and in rats, enhances trophoblast-guided spiral arteriole remodeling (7). Trophoblast signals leukocytes as well. In rats, uNKs are a target for trophoblast prolactin-like protein A (38). Trophoblast can act as a chemoattractant for macrophages (39) and can also induce vascular endothelium to secrete factors that attract uNKs and macrophages in a human co-culture model (40). In short, the cell types that appear key to vessel remodeling, trophoblast, uNK cells and macrophages, exist in complex relationships and their respective roles in spiral arteriole remodeling has yet to be fully elucidated.

The rat is a promising model system to use to better understand these processes. Like mice, rats have a rapid generation time and the capacity for powerful genetic manipulations (7,37,41–43). Like humans, rats have hemochorial placentation and a highly invasive trophoblast that reaches through the decidua and into blood vessels within the myometrium. In mouse, trophoblast invasion is shallow, predominantly interstitial rather than endovascular, and restricted to the mesometrial decidua (44,45). There are some key differences between the human and rat, to be sure. Rats have a bicornuate uterus and abembryonic implantation (46), unlike humans. Perhaps most notably, the rat’s deep placental bed is formed in a crescent-shaped split between the inner circular and outer longitudinal layers of uterine muscle. This space, which forms only in pregnancy, was originally called the metrial “gland” (47) or metrial triangle and it is the rat’s uterine-placental interface (48). This area is rich in immune cells and the tortuous spiral arterioles that serve the placenta. Despite differences between humans and rats, what we know of trophoblast-dependent remodeling appears conserved with humans (9,26) and the rat offers a model system in which we can interrogate the timing of remodeling events *in vivo* within the context of the entire implantation site to ultimately better understand the cellular mechanisms responsible.

Work in the rat on spiral arteriole remodeling has been confined almost exclusively to the later trophoblast-dependent phase of remodeling from mid to late gestation (13,31), typically from day 15 to 18 of the 23 day gestation period. Earlier decidua-associated remodeling is understudied, and we lack information on this earlier phase from the time the metrial triangle first appears to the initial invasion of the enlarged spiral arterioles by endovascular trophoblast. In this study, we present the rat as a tractable model system to investigate early remodeling of spiral arterioles in the uterine-placental interface. First we define the temporal window of early vessel remodeling in the rat between metrial triangle formation and the endovascular trophoblast invasion of parental arterioles. Second, we classify the vessels in the uterine-placental interface by stage of remodeling within that temporal window, based primarily on staining for smooth muscle actin. Third, we consider these events within the context of leukocyte and trophoblast invasion of the placental bed. With this framework, we present the rat as a model system that can be used to interrogate the process of early phase spiral arteriole remodeling.

## Methods

### Animals

Sprague-Dawley CD rats (*Rattus norvegicus*) were purchased from Charles River Laboratories (Wilmington, MA) between 2-3 months of age and maintained on a 10hr light:14hr dark cycle with water and food (Teklad 18% protein rodent diet) available *ad libidum*. All experimental protocols were performed in accordance with the National Resource Council’s <u>Guide for the Care and Use of Laboratory Animals</u> (49) and were approved by the Mount Holyoke College IACUC (BR-58-0613 and BR-63-0622). Virgin females were housed in pairs and their estrous cycles followed by daily vaginal lavage (50). Mating pairs were set up within one hour of lights out on the day of the female’s proestrus in a 2’ x 4’ enclosure with a clear Plexiglas front and blacked-out back and sides (FLN-MAR, Holyoke, MA). Mating was confirmed by vaginal lavage performed late in the following light cycle (the next morning). The morning that sperm were found in a vaginal lavage was considered day 1 of pregnancy. Pregnant rats were housed singly for the duration of gestation then sacrificed by CO2 inhalation and thoracotomy. The sample size was four dams on each of gestational days 9, 10, 12, and 14, two dams on each of days 11 and 13, and three dams on gestation day 16.

### Tissue preparation Classical Histology

Sections of uterine horn were pinned out in a dissecting tray, moistened with neutral buffered formalin to prevent curling, and individual segments of the uterine horn with some mesometrium still attached were preserved in Neutral Buffered Formalin (NBF). The pronounced size difference between earlier and later stage implantation sites made slightly different dissection and fixation protocols necessary. Smaller gestation day 9-12 samples were placed directly into NBF and held there for 24 hours prior to transfer to 70% ethanol. The larger gestation days 13-16 samples were punctured antimesometrially to let fluid escape prior to fixation, making the overall size of the tissue sample smaller while preserving the relative positions of the placenta, metrial triangle, and layers of uterine muscle. These larger samples were held in NBF for 48 hours before transfer to 70% ethanol. Following paraffin infiltration (Pioneer Valley Life Sciences Institute, Springfield MA) and embedding the entire implantation site was cut in 7um serial sections perpendicular to the long axis of the uterine horn, bisecting the chorionic plate and capturing the layers of uterine muscle and the uterine-placental interface in cross-section. The thirty or so serial sections closest to the center of the implantation site were mounted and stained with Hematoxylin & Eosin or with Masson’s Trichrome.

### Immunohistochemistry

Sections of uterine horn were pinned out in a dissecting tray and moistened with saline. For some samples, the uterus was moistened with 4% paraformaldehyde to prevent curling. These samples overfixed and were excluded from analysis. Segments of uterine horn were embedded in Tissue-Tek OCT Compound (Sakura Finetek) in an isopentane bath held over liquid nitrogen, then stored as OCT blocks at −80°C until cutting on a cryostat (−8°C) into 7um sections. The entire center portion of the implantation site was sectioned and mounted, and then the serial sections flanking the center were identified retrospectively and saved. Sections were taken through the entire implantation site, perpendicular to the long axis of the uterus, to capture the placenta, invading trophoblast cone, and metrial triangle in cross section.

Frozen mounted tissue sections were fixed with ice-cold acetone and run through a standard indirect immunoperoxidase staining protocol as follows. Endogenous peroxidase activity was quenched with 0.3% peroxide, nonspecific binding was blocked with 10% normal goat serum (Jackson Immunoresearch, West Grove, PA), and then sections were incubated with the primary antibody overnight at 4°C. The primary antibodies were: ASM-1, mouse anti smooth muscle actin (Millipore) to detect smooth muscle and used at a dilution of 1:1000; C-11, mouse anti-pan cytokeratin (Sigma) to detect trophoblast and used at a dilution of 1:10,000; PECAM-1 (clone E-4), mouse anti-CD31 (Santa Cruz Biotechnology) to detect endothelial cells and used at a dilution of 1:500; and OX-1, mouse anti rat CD45 (BD Pharmingen) to detect leukocytes and used at a dilution of 1:100. Negative control sections (one per slide) were incubated with Tris buffered saline. After overnight incubation with the primary antibody/TBS, sections were incubated for at least one hour at room temperature with a goat anti-mouse biotin-SP conjugated secondary antibody followed by peroxidase-conjugated streptavidin (2.2ug/ml and 1.5ug/ml, respectively, Jackson Immunoresearch). Slides were washed in TBS/0.1% Tween 20 between each step. To elicit the enzymatic color reaction we used an AEC substrate kit (Vector Laboratories) then lightly counterstained with Gill’s #1 hematoxylin so as not to interfere with ImageJ analysis of the antibody staining. Immunohistochemistry was performed on a single implantation site from dams who carried pregnancies to gestation day 9, day 10, day 11, day 12, day 13, day 14 and day 16, with sample sizes as indicated earlier.

### Quantification

Tissue sections capturing the entire uterine-placental interface of the rat and stained with an antibody against alpha smooth muscle actin, ASM-1 (Millipore), were photographed on an Olympus BX51 microscope using CellSens Basic software. The entire interface or metrial triangle area, defined as follows, was photographed. The uterine myometrium is composed of an inner layer of circular muscle and an outer layer of longitudinal muscle. The metrial triangle develops in a split between the two layers, forming a crescent shape whose thickest portion is typically centered over the tip of the invading cone of trophoblast. The curved base of the metrial triangle borders the uterine decidua and is perforated in midpregnancy by invasive trophoblast. We photographed both the thickest, vascularized center of the crescent and the tapered tips where the two layers of myometrium come back together. Using the “stitch” plugin within ImageJ (51) we then generated a composite image of the entire metrial triangle from individual images photographed at 100x. This enabled us to individually identify and number the blood vessels in each embryo’s uterine-placental interface.

Individual color photomicrographs were rendered into segmented binary images using ImageJ. Using the “threshold” function, we extracted the ASM-1 positive areas out from any background staining and took measurements from binary images in which ASM-1 positive areas stood out as white rings or flecks on a black background. Veins were excluded on morphological grounds, primarily their very thin and uniform vascular smooth muscle media but also a frequently squashed appearance in cross section. Vessel measurements were performed on the segmented binary images, except where a vessel showed negligible ASM-1 immunostaining. In those cases the vessel lumen area and diameter were measured on the original color photomicrograph. Immunostained vessels were measured from dams who carried pregnancies to gestation day 9, day 10, day 12, day 14 and day 16. Vessels from the complete metrial triangle area of a single embryo’s implantation site per dam were measured.

## Results

### In rat, trophoblast invasion of the spiral arterioles is deep in the metrial triangle

The uterine-placental interface in the rat is a crescent-shaped space between the inner and outer layers of the uterine muscle, a space bordered on its curved base by the mesometrial decidua, and topped near its peak by the point of entry for the radial arteries (Figure 1A, dotted line in inset photo). The branching uterine arteries in this zone are invaded by trophoblast, shown here on gestation day 13 (Fig. 1A, bluish purple stream of trophoblast perforates the base of the metrial triangle and reaches uterine blood vessels). Once tapped by trophoblast, blood in the arterioles flows toward the placenta in a central canal within the column of invading trophoblast and reaches the placenta’s transfer zone, the labyrinth. Figure 1B shows the dark purple trophoblast reaching enlarged blood vessels within the uterine-placental interface, and in 1C (arrow) the enlarged purple trophoblast cells are visible forming part of the wall of the blood vessel.

**Figure 1.**
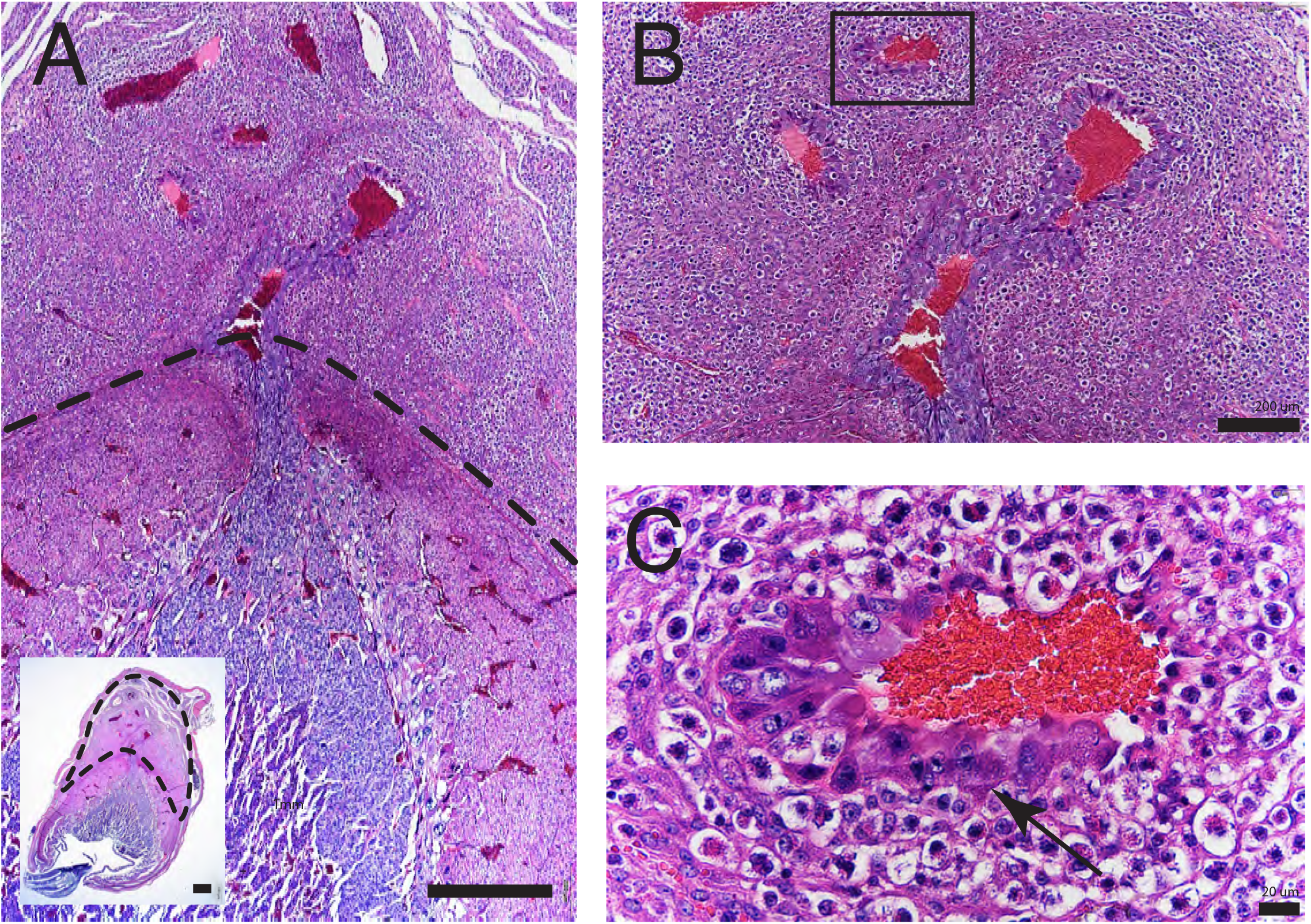
Rat placentation is deeply invasive and the metrial triangle is the vascularized zone where invasive trophoblast taps into the uterine arterial tree. The triangular compartment forms in early pregnancy and is the point of entry (from the parental side, top) for the vessels of the uterine arterial tree. Metrial triangle outlined with dotted black line, panel A inset. Base of metrial triangle, penetrated by bluish trophoblast of the central canal (from the fetal side, bottom), shown with dotted line in main image of panel A. Shown here on gestation day 13, fetally-derived trophoblast has entered the compartment and tapped into the vessels there, initiating the perfusion of the placental transfer zone, the labyrinth, with parental blood. Panels B and C, closeups of vessels in the metrial triangle. Vessel shown in panel C indicated by box in panel B, and lined along bottom and left edge with large, purplish endovascular trophoblast cells (arrow). Scale bars: panel A main and inset scale bars 1mm; panel B scale bar 200um; panel C scale bar 20um.

The rat’s uterine-placental interface zone becomes larger and more vascularized as pregnancy progresses. The interface enlarges in cross-sectional area between days 9 and 16 of pregnancy (Figure 2A) and the number of vessels in the space increases in the same time period (Fig. 2B). Not only does the overall area increase, but the proportion of interface occupied by blood vessels, a measure of vascularity, increases as well. Summing the lumenal area of all the vessels in a single metrial triangle, and dividing it by the overall area of that metrial triangle, shows that the overall perfusion of the uterine-placental interface by maternal blood increases significantly between days 9 and 16 of gestation in the rat (Fig. 2C).

**Figure 2A.**
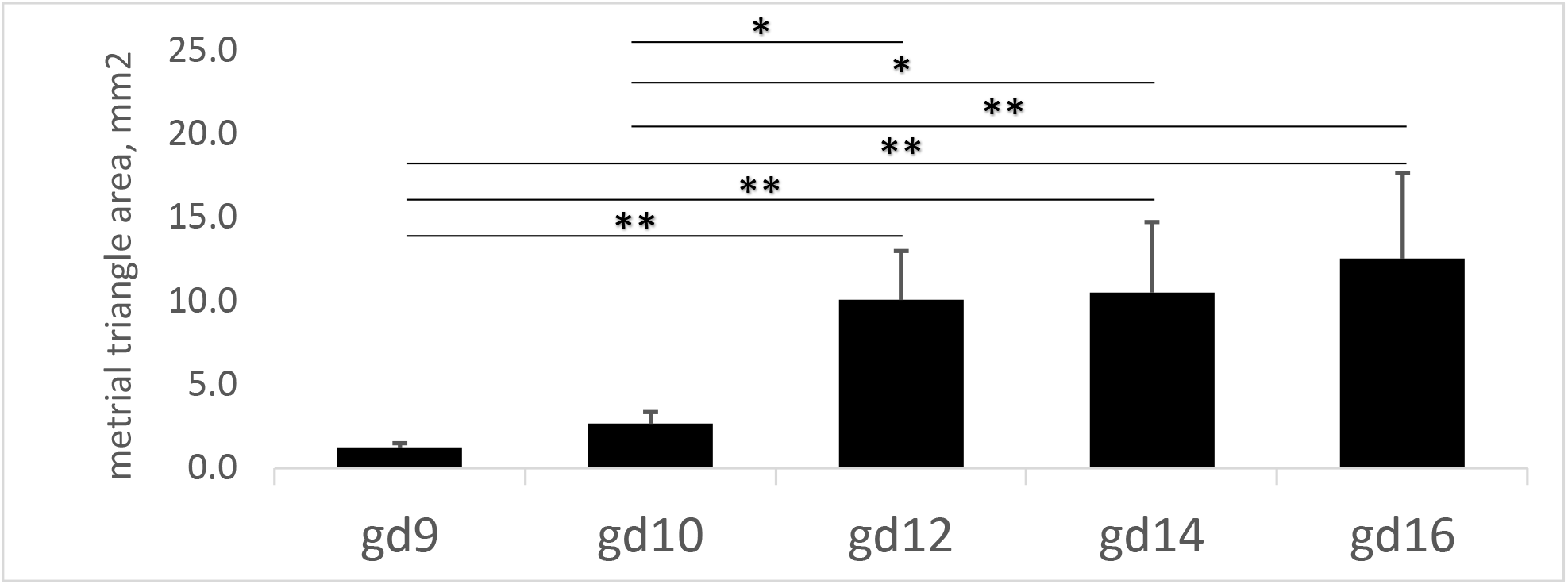
The area of the uterine-placental interface increased between gestation day 9 and 16. Whole metrial triangles were photographed at 100x, then individual images were stitched into a composite within ImageJ (51) and measured. *p<.05, **p<.01 by ANOVA with Tukey post-hoc comparisons.

**Figure 2B.**
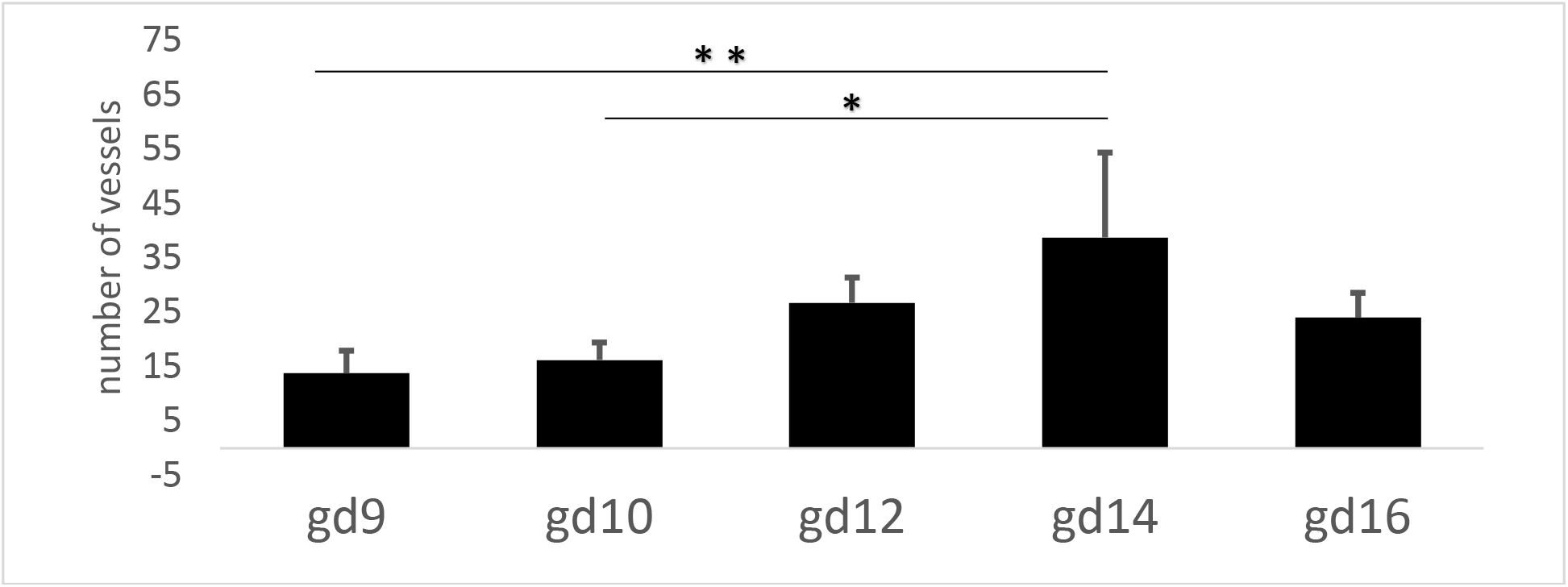
The number of vessels in the uterine-placental interface’s arterial tree increased from gestation day 9 to 14. *p<.05, **p<.01 by ANOVA with Tukey post-hoc comparisons.

**Figure 2C.**
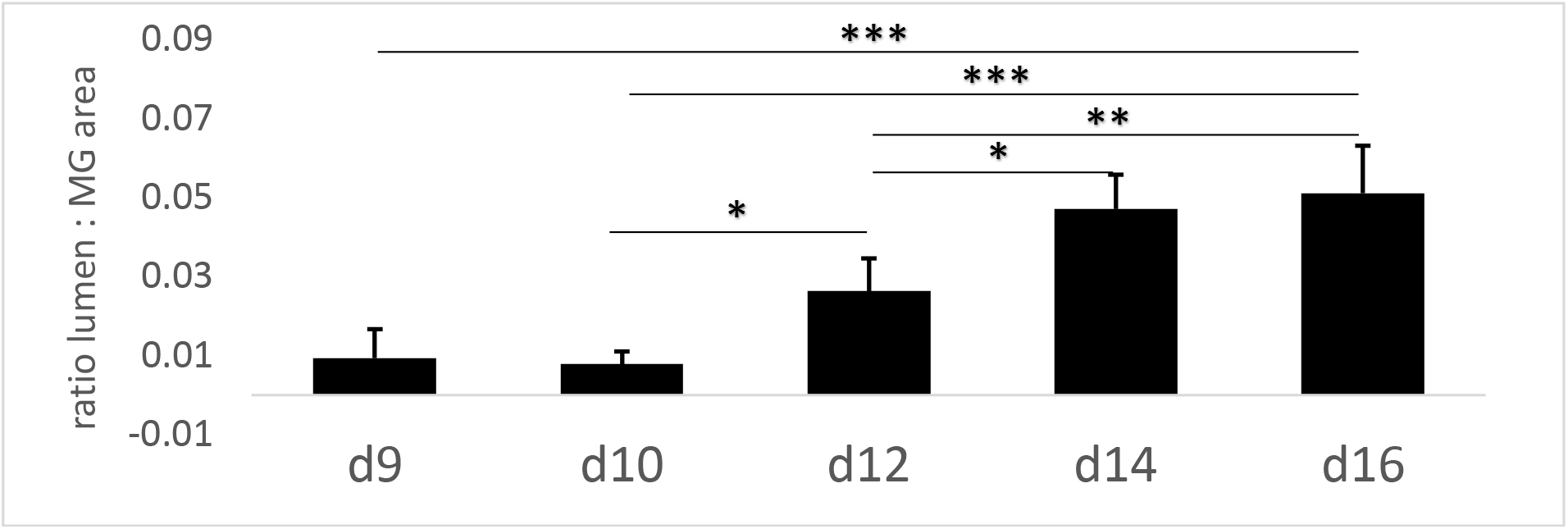
The vascularity of the uterine-placental interface increased between gestation days 9 and 16. Lumenal area of all vessels in the arterial tree were summed for each metrial triangle, then divided by the overall area of that metrial triangle in um^2^. *p < .05, **p< .01, ***p < .001 by ANOVA with Tukey post-hoc comparisons.

### Spiral arterioles are modified shortly after the uterine-placental interface is established

Initially, the arterioles within the metrial triangle are small, with a pink (H&E) or red (Masson’s Trichrome) intact smooth muscle media layer (see four small vessels in Figure 3A and 3B). Even at this early stage (gestation day 10 shown) the smooth muscle nuclei in the media layer appeared jumbled, and the endothelial cells often protruded into the lumen as is visible in Fig. 3 panels A-D.

**Figure 3:**
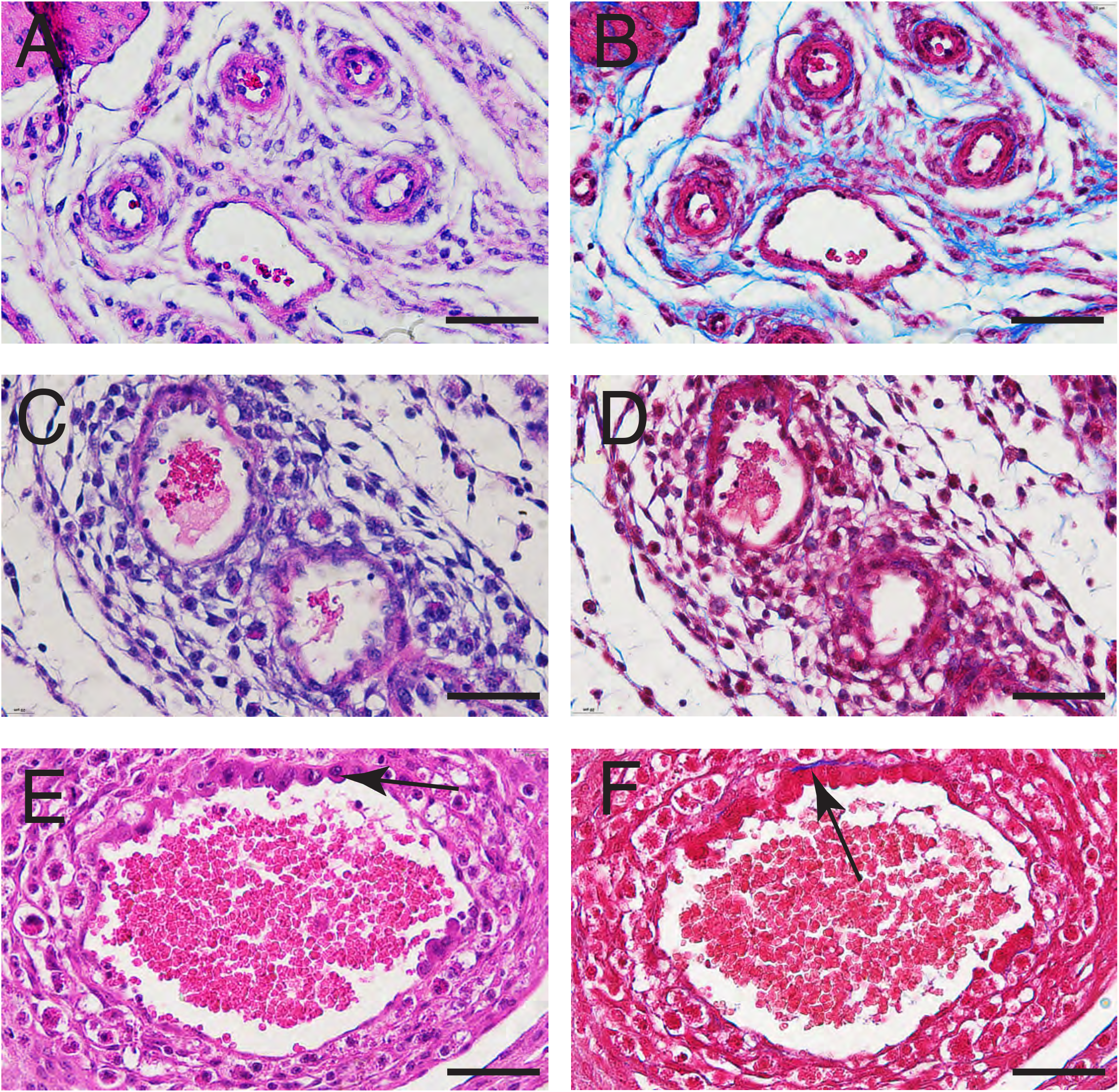
Arterioles in the uterine-placental interface zone are progressively enlarged into high-flow, low pressure conduits carrying parental blood to the placenta’s transfer zone. Tissues of the uterine-placental interface were stained with H&E (A, C, E) and Masson’s Trichrome (B, D, F). Shown are intact vessels on gestation day 9 (A,B), enlarged vessels with thinned and irregular smooth muscle media layer on day 12 (C, D) and very dilated vessels on day 16 (E, F) partially lined with trophoblast. Arrow (E) indicates endovascular trophoblast. Arrow (F) indicates fibrinoid in blue. Scale bars 50um

The vessels enlarge as the smooth muscle thins and develops small breaks (Figure 3C and 3D). Ultimately, the vessel dilates to many times its original size and invasive endovascular trophoblast (enlarged cells along top border of vessel in 3E (arrow)) replaces the smooth muscle. Fibrinoid (blue in Masson’s Trichrome staining, Fig. 3 panels B, D, and F) was very thinly spread around the smaller vessels, and portions of the vessel wall only showed any fibrinoid where trophoblast had replaced the media layer and the vessel in question was also close to the basal portion of the metrial triangle. An example of a distinct fibrinoid deposit along the basal surface of the endovascular trophoblast is shown in Figure 3F (arrow). The scarcity of fibrinoid is likely because the study window of day 9 to day 16 ends before many vessels in the metrial triangle have been lined with endovascular trophoblast.

The progressive dilation of vessels is a factor of changes to the arterioles’ vascular tunica media layer. We set out to categorize the stages of vessel deconstruction by measuring the changes to the arteriole smooth muscle media layer. Using tissue immunostained for smooth muscle (the primary antibody ASM-1) blood vessels (n=442) were categorized by the percent of the lumen surrounded by muscle. Vessels that were classified as 100% intact had a smooth muscle media surrounding the lumen; vessels that were 0% intact had lost their smooth muscle entirely. Figure 4 shows that on each day of the study period, the vessels’ vascular smooth muscle media was in a variety of phases of deconstruction. On gestation days 9 and 10, most spiral arterioles were completely intact, but a small number of vessels (less than 10%) showed small breaks. Gestation day 12 was the first day that more advanced fragmentation of the tunica media became evident. The proportion of vessels showing extensive fragmentation steadily increased through gestation day 16. Spiral arterioles in the metrial triangle that had no vascular smooth muscle at all were visible on days 14 and 16.

**Figure 4.**
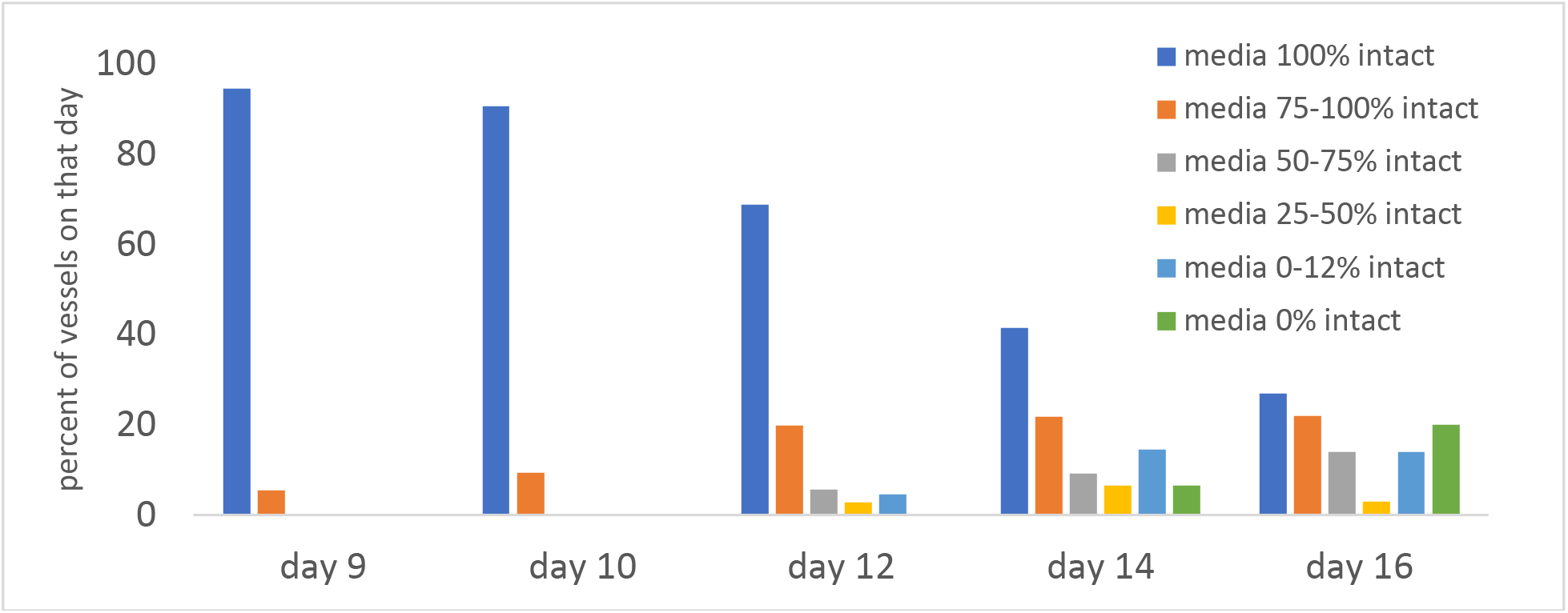
The extent of vascular smooth muscle fragmentation increased with gestational age, and vessels were in multiple degrees of remodeling on each gestational day. Tissue samples stained with the anti-smooth muscle antibody ASM-1 were photographed and rendered as binary black and white images to highlight the areas of positive staining, then measured in ImageJ. Vessels in these binary images were categorized based on the percent of the vessel wall that was immunopositive for ASM-1, an estimate of the percent of the media layer that remained intact. n = 442 blood vessels from 19 implantation sites on days 9 (n=56), 10 (n=64), 12 (n=106), 14 (n=152), and 16 (n=64) of pregnancy.

The primary variable used to break the remodeling process into distinct phases was the extent of the vessel media that remained intact (Table 1). Categorizing vessels by media “intactness” was a reasonable predictor of other progressive changes to vessel morphology. In particular, intactness predicted vessel media thickness as well as lumenal diameter and area. Vessels split into four groups based on the degree of media fragmentation showed that as the vascular smooth muscle fragmented, it progressively thinned (Table 1, between groups ANOVA p<.001). Stages based on the degree of media fragmentation also showed a progressive increase in vessel diameter (p<.001), and the average area of the vessel lumen (p<.001).

**Table 1.** Summary of the stages of early vessel remodeling in the rat uterine-placental interface

| Stage | 1 | 2 | 3 | 4 |
| --- | --- | --- | --- | --- |
| media intactness (ASM-1) | (a) 100% intact, sometimes (b) intact but vacuolated or wispy | small breaks (at least 75% intact) | 25-75% intact | disorganized fragments (25% or less intact) |
| media thickness, $\mu\text{m}$ | $8.8 \pm 3.5 \mu\text{m}$ | $7.4 \pm 3.1$ | $5.4 \pm 3.0$ | $1.4 \pm 2.6$ |
| vessel diameter, $\mu\text{m}$ | $44.2 \pm 35.4 \mu\text{m}$ | $106.4 \pm 82.3$ | $135.2 \pm 75.3$ | $225.9 \pm 123$ |
| vessel lumen area, $\mu\text{m}^2$ | $2102.2 \pm 4363.4 \mu\text{m}^2$ | $11532.5 \pm 20214.8$ | $16782.0 \pm 15375.1$ | $49053.3 \pm 50308.1$ |
| vessels measured | n = 263 | n = 76 | n = 43 | n = 56 |
| endothelium (PECAM-1) | (a) intact, (b) swollen | fragmented | fragmented or undetectable | occassional PECAM-1 staining |
| endovascular trophoblast (C-11) | Absent | Absent | Absent | Present in vessels near base of metrial triangle from gestation day 13 on |
| fibrinoid (Masson's Trichrome) | Undetectable or thinly surrounding media | Undetectable or thinly surrounding media | Undetectable or thinly surrounding media | Present in large vessels with endovascular trophoblast from gestation day 13 on |

Notably, vessels categorized as *intact* were not necessarily *unmodified*. Vessels in Stage 1, completely surrounded by smooth muscle, were a heterogenous group. A majority of vessels in this stage had jumbled smooth muscle nuclei in the media layer, and many had endothelial cells protruding into the lumen. Some ASM-1 immunostaining suggested that the vascular smooth muscle was vacuolated in Stage 1. In other cases, the ASM-1 staining could appear “wispy” and light, suggesting that the muscle cells may have been freed from their matrix in the media layer. Such intact but modified vessels were present even as early as gestation day nine.

Staining for CD31 with PECAM-1 showed that Stage 1 vessels had an intact endothelium, sometimes with the cells protruding into the lumen (Table 1). Endothelium fragmented around the same time that breaks first appeared in the vascular smooth muscle (Stage 2). As the smooth muscle became increasingly disrupted, CD31/PECAM-1 immunostaining became fainter and at times difficult to distinguish from background.

The vessel smooth muscle was very fragmented before any endovascular trophoblast was visible lining the lumen. Media disorganization and fragmentation preceded the invasion of any endovascular trophoblast, which became visible as of day 13. There was minimal evidence of fibrinoid deposition in vessels partially lined with trophoblast. Vessels with a substantial amount of trophoblast in their wall were temporally restricted to the days 13-16 in this study, and anatomically restricted to the basal portion of the metrial triangle, close to the developing central canal as it emerged from the decidua.

Figure 5 shows immunohistochemical staining within the metrial triangle to back up the data presented in Table 1. Initially, smooth muscle completely encircled the arteriole lumen (Figure 5A) and became irregular, suggesting disorganization (Fig. 5D) before fragmenting (Fig. 5G, 5J). In unmodified vessels with an intact media, CD31/PECAM-1 staining for endothelium shows a thin, intact endothelium lining the arteriole (Fig. 5B) that appeared to swell and expand (Fig. 5E) concurrent with the initial disorganization of the smooth muscle. As the vascular smooth muscle fragmented (Fig. 5G and 5J), PECAM-1 staining for endothelium also showed some fragmentation (Fig. 5H, 5K). Trophoblast immunostaining (C-11) was absent from the vessel walls when the media remained intact in the earliest phases of media fragmentation (Fig. 5C, 5F), and endovascular trophoblast was visible in the metrial triangle lining only those vessels with very fragmented or absent smooth muscle (Fig. 5L). Trophoblast was only visible in the metrial triangle on or after day 13 of gestation.

**Figure 5.**
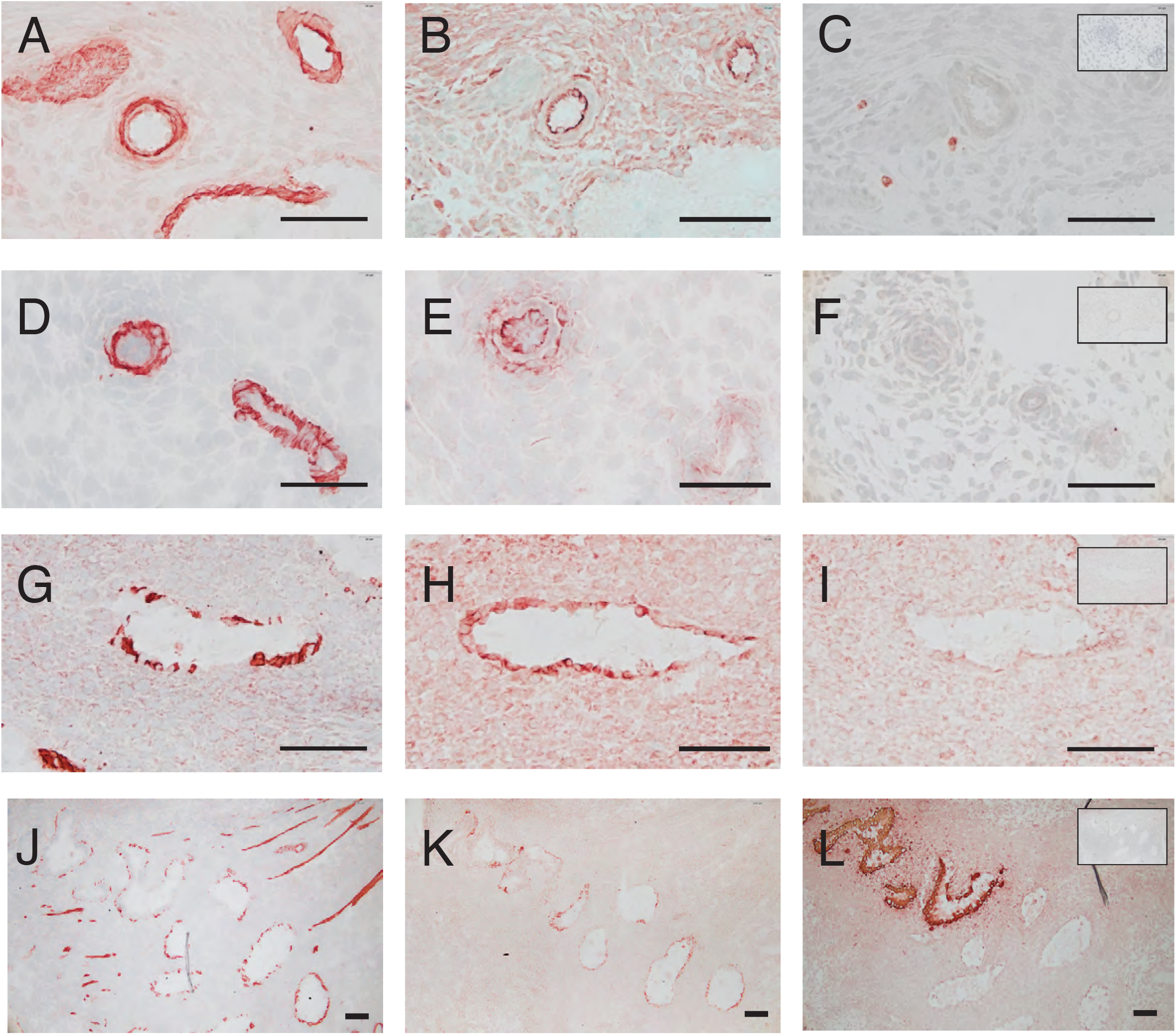
In rat, early disruption of smooth muscle and endothelium is followed by trophoblast invasion into vessels with a very fragmented or absent smooth muscle media. Top row (panels A, B, C) images taken of gestation day 9 metrial triangle at 500x; second row (D, E, F) gestation day 10 at 500x; Third row (panels G, H, I) gestation day 13 at 500x; fourth row (panels J, K, L) images taken of gestation day 14 metrial triangle at 100x. The first column (panels A, D, G, J) shows antibody staining for smooth muscle actin with ASM-1; tissues in the second column (panels B, E, H, K) are stained with antibody to CD31/endothelium, PECAM-1; the third column (panels C, F, I, L) shows tissues stained with antibody to cytokeratin, C-11, for trophoblast. Negative controls (TBS in lieu of primary antibody) shown as insets in third column. All scale bars 100um.

Vessels at multiple stages of deconstruction were found on all days in this study. Remodeling was not synchronous within a particular metrial triangle or a particular gestational day. Even so, the appearance of immune cells and trophoblast, known in other species to be key players in remodeling, was gestation-day dependent. An infiltration of immune cells into the metrial triangle was visible on gestation day 10 (Figure 6B) at a time that the vessel media were intact (Fig. 6A) and trophoblast (Fig. 6C) was still restricted to the ectoplacental cone overlying the embryo deep in the decidua. CD45/OX-1 positive cells very densely clustered around vessels in the metrial triangle on day 12 (Fig 6E). This pronounced infiltration of immune cells was evident on the first day (day 12) that vessels began to show extensive fragmentation of the arterioles’ tunica media (Figure 4). Trophoblast was first visible in the metrial triangle on gestation day 13 (Fig 6 C, F, I).

**Figure 6.**
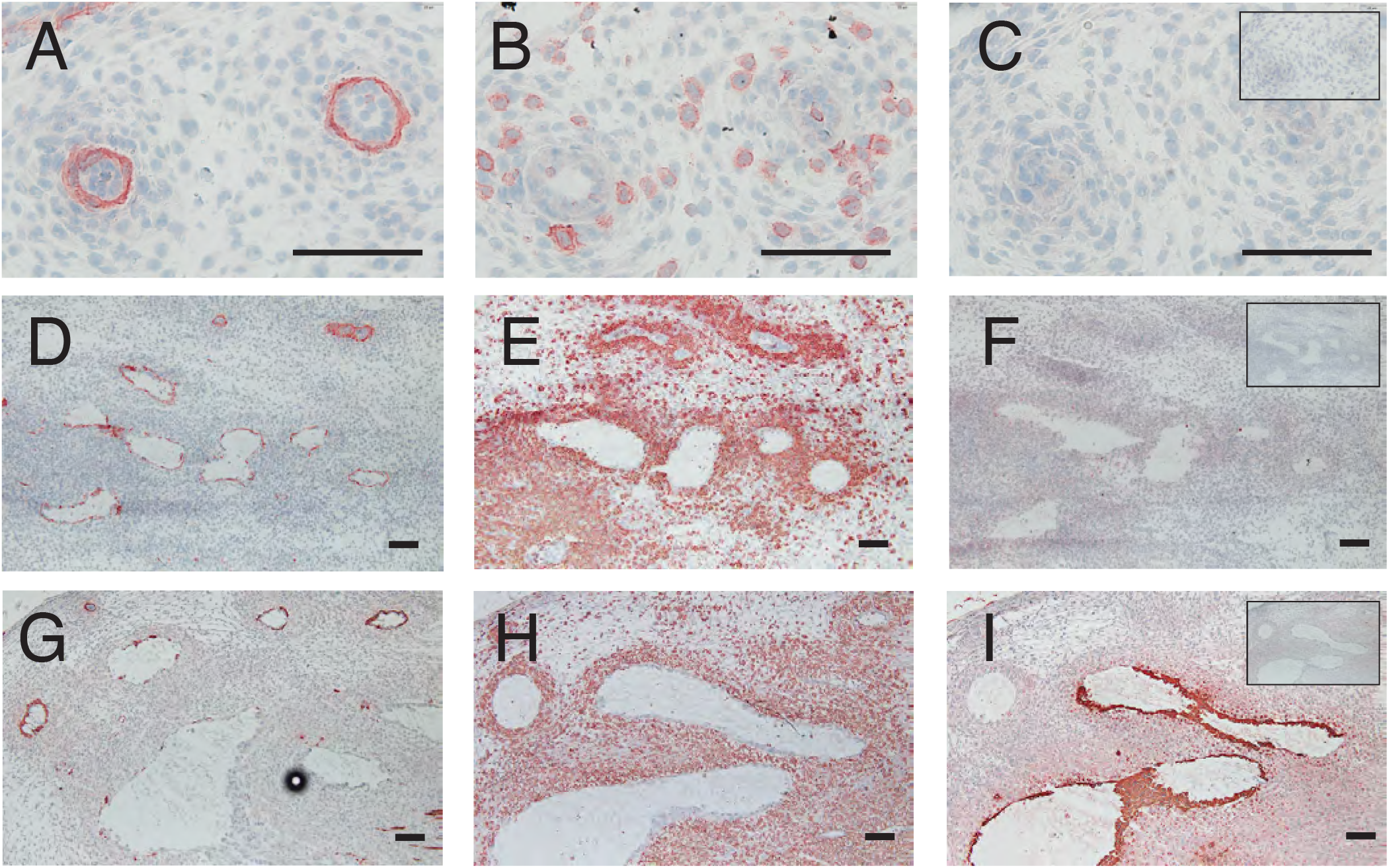
While vessel remodeling is not synchronous with day of gestation, the appearance of immune cells and trophoblast in the metrial triangle is dependent on day of gestation. Shown are gestation day 10 (A, B, C) at 500x; day 12 (D, E, F) at 100c and day 14 (G, H, I) at 100x. Tissues in the left-hand column (A, D, G) antibody stained with ASM-1 for smooth muscle; center column (B, E, H) with CD45/OX-1 for immune cells; and right column (C, F, I) stained for trophoblast with C-11. Negative controls (TBS in lieu of primary antibody) shown as insets in right hand column. All scale bars 100um.

## Discussion

To establish the rat as a model system for understanding early decidua-associated remodeling of spiral arterioles in the deep placental bed, in this study we describe the architecture of the metrial triangle and the early changes in the spiral arterioles of the rat’s uterine-placental interface prior to the full replacement of the vessel wall by invasive trophoblast. The study period opened shortly after the metrial triangle area first formed in the rat (52). The study period ended as trophoblast partially lined the largest, most fragmented vessels at the base of the metrial triangle on gestation day 16. We identify several temporal landmarks in this period that could be fruitful for investigating the mechanisms of this early phase spiral arteriole remodeling.

The uterine-placental interface increased in size and vascularity during this time. Vascularity, the proportion of the metrial triangle composed of vessel lumena, began to increase prior to endovascular trophoblast invasion. In this period, blood in the uterus is redirected from the decidua basalis to the chorioallantoic placenta in rats (25). The rapid and dramatic expansion of the blood supply to the placenta in this period underscores the importance of this zone to securing an oxygen supply to support rapid growth and increasing metabolic demands of the mammalian embryo (53). The vascularization, and concomitant oxygen delivery, influence lineage determination in the very trophoblast that will invade days later (7).

There was a progressive deconstruction of the spiral arterioles evident as early as day 9, well before trophoblast invasion of the metrial triangle. Vessels at multiple stages of remodeling were visible on every day of gestation from days 9 to 16. Unlike embryo development, which can be precisely staged by day of gestation, the progressive disassembly of the spiral arterioles was not synchronized with gestation day. For this reason we felt that a staging system that used gestation day as a proxy for stage of remodeling, as Charalambous et al. have done for decidua-associated remodeling in mice (30), might have masked the progression of changes.

Thus we divided the continuous remodeling process into stages based primarily on the morphology of the vascular smooth muscle as identified by ASM-1 immunoreactivity (Table 1). Using degree of media “intactness” as the primary staging criterion was fortuitous in that changes in vessel shape and cellular composition correlated well with the degree of smooth muscle fragmentation. For example, vessels with an intact and apparently organized vascular smooth muscle (Stage 1a) had a thin, intact endothelium while vacuolated or disorganized vascular smooth muscle media seemed correlated with swollen endothelial cells protruding into the vessel lumen (Stage 1b). In addition, the first small breaks appearing in the vascular smooth muscle layer and the first small breaks in the endothelium appeared coetaneously (Stage 2). In the wide range of 25% to 75% media intactness, further changes to the endothelium were not evident, so it seemed reasonable to combine this wide range into a single category (Stage 3). Endovascular trophoblast was visible exclusively in those vessels with very fragmented or absent vascular smooth muscle (Stage 4). Its presence in the vessel was a product of both day of gestation (day 13 or later) and vessel placement within the metrial triangle. Spiral arterioles with very fragmented muscle located close to the base of the metrial triangle were the most likely to contain trophoblast and those located peripherally were less likely to contain trophoblast. Trophoblast invasion of the metrial triangle typically begins near the midpoint of the triangle’s base, and once it perforates the circular muscle forming the boundary with the decidua, trophoblast spreads from the central zone outward (13,31,54).

The steps in spiral arteriole remodeling seen in this study are largely consistent with what has been reported in humans by Smith et al. (2006) (28) and by Pijnenborg et al. (2006) (55): first, the disruption and partial loss of the vascular smooth muscle, with endothelial vacuolation (22). Next the muscle media layer begins to break up, and sometimes interstitial trophoblast is visible outside the vessel at that time (55). As the endovascular trophoblast migrates into the vessel, the endothelium begins to show breaks and fibrinoid appears underneath the endovascular trophoblast beginning around day 15 and increasing through days 16 and 17 (9). Full transformation is achieved when the vessel becomes sac-like with a wall of endovascular trophoblast embedded in fibrinoid.

In the rat it appears that early vessel remodeling may already be underway on gestation days 9 and 10. While the canonical image of an arteriole shows spindle-shaped nuclei and a very smooth, regular media layer, that was not consistently reflected in our samples. Rather, the vascular smooth muscle in vessels appeared lumpy and irregular and its nuclei were often rounded and jumbled-looking. Vessel media disorganization may begin shortly after the metrial triangle forms in rat. Charalambous et al. (30) working in mice observed the muscular wall of the spiral arteriole thickening even prior to uNK infiltration. In human tissue, Smith et al. (24) note a kind of “loosening” of the muscle with the degradation of collagen type IV in the vascular smooth muscle also prior to uNK infiltration.

Small numbers of immune cells were present around the vasculature in day 10 samples, and in gestation day 12 tissues, CD45/OX-1 positive cells were visible densely clustered around the arterioles in the metrial triangle. On that day arterioles were at a wide variety of different phases of remodeling. Many of the OX-1 immunoreactive cells were likely uNKs, which are and prominent in the metrial triangle in rodents (56,57) as well as abundant in human uterus (58). Macrophages are also present in the uterus throughout pregnancy (59) though without the same peak density as uNKs during remodeling. Both uNKs and macrophages have been found embedded in the wall of remodeling vessels in human *in vitro* coculture systems (60), expressing MMPs (29) and producing secretory products capable of inducing muscle cell rounding and separation (32). Leukocyte infiltration has also been associated with endothelial swelling (28). In mice, uNKs infiltrate the vessel wall as the media layer progressively disorganizes (30).

Spiral arteriole remodeling is typically described as occurring in two phases, an early “decidua-associated” or “uNK-associated” phase and a later “trophoblast dependent” phase. But recent work in rats (21) complicates the idea of leukocyte-associated remodeling that is independent of trophoblast. The vascular smooth muscle media can contain cells of different phenotypes (61), including contractile muscle cells and synthetic smooth muscle cells that are involved with vessel maintenance and remodeling. Nandy et al (2020) showed that vascular smooth muscle retained its contractile phenotype even when uNKs were present in the perivascular space. It was the presence of invasive trophoblast surrounding the arterioles that was correlated with a switch in smooth muscle phenotype from contractile to synthetic. In the present study, our use of the ASM-1 antibody against alpha actin may well have missed muscle cells of the synthetic, non-contractile phenotype. In future being able to distinguish the different types of smooth muscle could help elucidate the cellular mechanisms behind the earliest remodeling events that involve disorganization of the smooth muscle media.

These results of this study show that in the rat there are significant changes to the spiral arterioles and the placental bed shortly after the metrial triangle forms. These changes include a significant increase in vascularity, very early disorganization and fragmentation of the smooth muscle media, and a progression of subsequent anatomical changes in spiral arterioles akin to the early changes in humans. In the rat these changes can be studied *in vivo* over time, making the rat a promising model system to better understand the complex relationships between cells in the unique microenvironment of the uterine-placental interface.

## Acknowledgements

The authors thank Jennifer Ser-Dolansky and Kelly Gregory, Ph.D., of the Pioneer Valley Life Sciences Institute for support with histology, Kathleen Byrne for maintaining the animals, and Heather Hamilton of the Mount Holyoke College Science Center Microscopy facility for support with brightfield microscopy. We also thank the Mount Holyoke College Department of Biological Sciences and the Mount Holyoke LYNK internship program for financial support to YZ. This research did not receive any specific grant from funding agencies in the public, commercial, or not-for-profit sectors.

